# The tolerance to hypoxia is defined by a time-sensitive response of the gene regulatory network in sea urchin embryos

**DOI:** 10.1101/2020.08.09.242933

**Authors:** Majed Layous, Lama Khalaily, Tsvia Gildor, Smadar Ben-Tabou de-Leon

**Author notes:** Correspondence: Smadar Ben-Tabou de-Leon.

## Abstract

Deoxygenation, the reduction of oxygen level in the oceans induced by global warming and anthropogenic disturbances, is a major threat to marine life. This change in oxygen level could be especially harmful to marine embryos that utilize endogenous hypoxia and redox gradients as morphogens during normal development. Here we show that the tolerance to hypoxic conditions changes between different developmental stages of the sea urchin embryo, due to the structure of the gene regulatory networks (GRNs). We demonstrate that during normal development, bone morphogenetic protein (BMP) pathway restricts the activity of the vascular endothelial growth factor (VEGF) pathway to two lateral domains and by that controls proper skeletal patterning. Hypoxia applied during early development strongly perturbs the activity of Nodal and BMP pathways that affect VEGF pathway, dorsal-ventral (DV) and skeletogenic patterning. These pathways are largely unaffected by hypoxia applied after DV axis formation. We propose that the use of redox and hypoxia as morphogens makes the sea urchin embryo highly sensitive to environmental hypoxia during early development, but the GRN structure provides higher tolerance to hypoxia at later stages.

**Summary statement:** The use of hypoxia and redox gradients as morphogens makes sea urchin early development sensitive to environmental hypoxia. This sensitivity decreases later, due to the structure of the gene regulatory network.

## Introduction

During the evolution of metazoans, animals were exposed to variations in oxygen levels and molecular mechanisms evolved to enable organisms to cope with hypoxic conditions (Semenza, 2012). However, it is still unclear whether these mechanisms are sufficient to protect marine organisms and specifically, their embryos, from the acute hypoxic conditions that become more common in the oceans (Altieri et al., 2017; Breitburg et al., 2018; Hughes et al., 2020). In the last 50 years the dissolved oxygen (O_2_) content of the global ocean has decreased by more than 2%, apparently due to warming that reduces oxygen solubility and increases biological consumption (Schmidtko et al., 2017). Recent studies indicate that oxygen loss in the oceans, termed deoxygenation, is more lethal to marine life than the direct effect of the rising temperatures or ocean acidification (Altieri et al., 2017; Breitburg et al., 2018; Hughes et al., 2020; Schmidtko et al., 2017; Vaquer-Sunyer and Duarte, 2008). The embryos of marine organisms could be highly sensitive to deoxygenation; especially embryos that use endogenous hypoxia and redox gradients as morphogens to guide the activation of gene regulatory networks (GRNs) during normal development (Chang et al., 2017; Coffman and Su, 2019; Cordeiro and Tanaka, 2020; Dunwoodie, 2009; Lendahl et al., 2009). Deciphering the structure and function of developmental GRNs that are activated by hypoxia and redox morphogens is a key to understand this fundamental regulatory mechanism as well as to assess the expected effect of ocean deoxygenation on marine embryos.

The sea urchin embryo provides an attractive system to study the developmental GRNs that are driven by variation in oxygen and redox levels and the effect of hypoxic conditions on these GRNs. Sea urchins are major grazers in shallow seas and coastal waters across the oceans (Pearse, 2006) and adult sea urchins were shown to be moderately sensitive to hypoxic conditions (Hughes et al., 2020; Low and Micheli, 2018; Suh et al., 2014; Vaquer-Sunyer and Duarte, 2008). The experimental advantages of sea urchin embryos and the role of the sea urchins in marine ecology make them a prominent model system for developmental and ecological studies (Pearse, 2006; Peter and Davidson, 2011; Sethi et al., 2012). The models of the gene regulatory networks that drive sea urchin early development are the state of the art in the field (Morgulis et al., 2019; Oliveri et al., 2008; Peter and Davidson, 2011). Importantly, the sea urchin GRNs use endogenous oxygen and redox gradients as developmental morphogens that drive the formation of the dorsal-ventral (DV) axis (Chang et al., 2017; Coffman et al., 2014; Suh et al., 2014).

During early development of the sea urchin embryo, maternally induced oxygen and redox gradients initiate the localized activity of several signaling pathways that eventually control the patterning along the DV axis (Fig. 1, (Chang et al., 2017; Coffman et al., 2014; Suh et al., 2014)).

**Figure 1.**
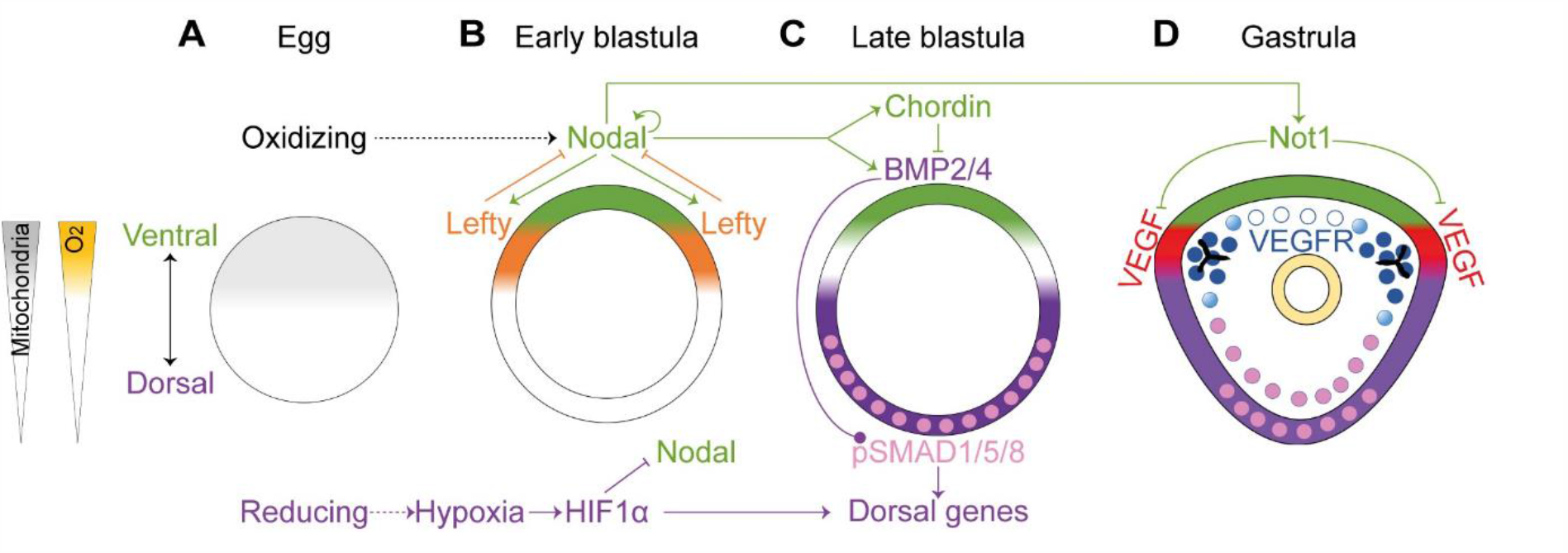
The regulation of DV axis formation downstream of redox and oxygen gradients in the sea urchin embryo. Diagrams showing sea urchin DV and skeletal patterning in developing sea urchin embryos in normal conditions based on (Chang et al., 2017; Coffman et al., 2009; Coffman and Davidson, 2001; Coffman et al., 2004; Coffman et al., 2014; Czihak, 1963; Duboc et al., 2004; Lapraz et al., 2009; Li et al., 2012). **(A)** The asymmetric distribution of mitochondria in the egg induces a redox gradient. **(B)** Regulatory interactions between *nodal, lefty* and HIF1α at the early blastula stage. **(C)** Nodal regulation of BMP signaling in the late blastula stage. **(D)** In the gastrula stage, Nodal activates the expression of Not1, that represses *VEGF* expression in the ventral ectoderm. Throughout the figure, the ventral side and Nodal expression domain are highlighted in green, the dorsal side and the domain of BMP activity are marked in purple. Nucleus that show pSMAD1/5/8 are highlighted in pink. *VEGF* expression is marked in red. *VEGFR* expression is marked in blue.

In the eggs of the sea urchins, the mitochondria are concentrated at the future ventral side (Coffman et al., 2004; Coffman et al., 2014), which leads to the formation of redox and oxygen gradients in the early embryos (Fig. 1A, (Agca et al., 2009; Coffman et al., 2004)). Specifically, the mitochondria produces reactive oxygen species (ROS) that generate an oxidizing environment which activate redox sensitive transcription factors that activate the expression of the Nodal ligand in the ventral ectoderm (Agca et al., 2009; Coffman et al., 2004; Coffman et al., 2014). Nodal reception drives the expression of the Nodal ligand and its antagonist Lefty and the positive and negative feedback interactions between these two proteins define the boundaries of the ventral ectoderm (Fig. 1B, (Duboc et al., 2008; Duboc et al., 2004)). Nodal activity drives the expression of the Bone Morphogenetic Protein (BMP), BMP2/4, and its antagonist Chordin, forming an incoherent feedforward loop (Fig. 1C, (Agca et al., 2009; Coffman et al., 2004; Coffman et al., 2014)). Chordin prevents the binding of BMP2/4 to its receptor at the ventral side so BMP is received only at the dorsal side where it activates gene expression through the phosphorylation of the transcription factor SMAD1/5/8 (Ben-Tabou de-Leon et al., 2013; Duboc et al., 2004; Lapraz et al., 2009) (Fig. 1C, D). Another early regulator of dorsal gene expression is the transcription factor, hypoxia-inducible factor 1α (HIF1α) that is stabilized in the dorsal side of the sea urchin blastula, apparently downstream of the oxygen gradient (Fig. 1A, B, (Ben-Tabou de-Leon et al., 2013; Chang et al., 2017; Coffman et al., 2009)). Thus, sea urchin embryos use the Nodal and BMP pathways and HIF1α to generate their DV axis downstream of redox and oxygen gradients inherited from the sea urchin egg.

Growth in hypoxic conditions leads to radialization of sea urchin embryos with prominent effects on the larval skeleton (Agca et al., 2009; Coffman et al., 2004). The skeleton of the sea urchin larvae is made of two skeletal calcite rods, the spicules, that are formed within a tubular syncytial chord produced by the skeletogenic cells (Morgulis et al., 2019; Oliveri et al., 2008). When the embryos are grown in hypoxic conditions the formation of the DV axis and the skeleton are disrupted, the spicules do not elongate properly and the embryonic morphology is significantly deformed (Agca et al., 2009; Coffman et al., 2004).

Sea urchin skeletogenesis depends on the Vascular Endothelial Growth Factor (VEGF) pathway, an essential regulator of vertebrates’ vascularization and of tubulogenesis in other phyla (Potente et al., 2011; Tettamanti et al., 2003; Tiozzo et al., 2008; Yoshida et al., 2010). The VEGF Receptor (VEGFR) is expressed in the sea urchin skeletogenic cells together with five transcription factors whose homologs are essential for vertebrates’ vascularization (Adomako-Ankomah and Ettensohn, 2013; Duloquin et al., 2007; Morgulis et al., 2019; Sun and Ettensohn, 2014). This and other similarities between the sea urchin skeletogenic gene regulatory network (GRN) and the vertebrates’ vascularization GRN suggest that these GRNs evolved from a common ancestral tubulogenesis GRN (Morgulis et al., 2019). The VEGF ligand is secreted from two lateral ectodermal domains located between the dorsal and the ventral ectoderm (Fig. 1D, (Adomako-Ankomah and Ettensohn, 2013; Duloquin et al., 2007; Morgulis et al., 2019)). VEGF expression is repressed in the ventral ectoderm by the transcription factor Not1 that is activated by Nodal signaling (Fig. 1D, (Li et al., 2012)). Yet, the regulatory links between BMP, HIF1α and VEGF signaling and how VEGF and BMP pathways are affected by hypoxia are not known.

Overall, sea urchin DV axis formation and skeletogenesis are strongly affected by hypoxic conditions and are regulated by Nodal, BMP, VEGF and HIF1, downstream of maternal oxygen and redox gradients (Fig. 1). To understand the effect of exogenous hypoxia on sea urchin development, here we study the regulatory links between the sea urchin DV and skeletogenic GRNs during normal development and under hypoxia applied at different developmental stages. We reveal that these two GRNs are strongly connected through the interactions between the BMP and VEGF pathways and that the DV GRN is hypersensitive to hypoxia during early development but becomes relatively tolerant to low oxygen levels with developmental progression.

## Results

### Sea urchin BMP2/4 controls skeletal patterning and VEGF expression

We first wanted to elucidate the links between BMP and VEGF signaling and the effect of BMP signaling on skeletogenic gene expression during normal sea urchin development. To that end, we knocked-down (KD) BMP2/4 expression by the injection of translation morpholino oligonucleotides (MO) into the eggs of the Mediterranean sea urchin species, *Paracentrotus lividus* (*P. lividus*, Fig. 2, see methods for details). Embryos injected with BMP2/4 MO show two major skeletogenic phenotypes: the formation of ectopic spicules in addition to the normal two spicules (ES, Fig. 2B) and ectopic skeletal branching, where the basic structure of two spicules is still observed (EB, Fig. 2C). These observations are in agreement with previous studies of BMP perturbations (Duboc et al., 2004; Lapraz et al., 2009). The expression level of *VEGF* is largely unchanged in BMP morphants (QPCR, Fig. 2G) but its spatial expression expands to one side of the ectoderm (detected by whole mount *in-situ* hybridization [WMISH], Fig. 2D). Since BMP signaling induces dorsal specification (Duboc et al., 2004; Lapraz et al., 2009), *VEGF* expansion in BMP morphants is most likely, to the domain that would normally be specified as dorsal ectoderm. Hence, this data implies that BMP activity represses *VEGF* expression in the dorsal ectoderm in normal embryos.

**Figure 2.**
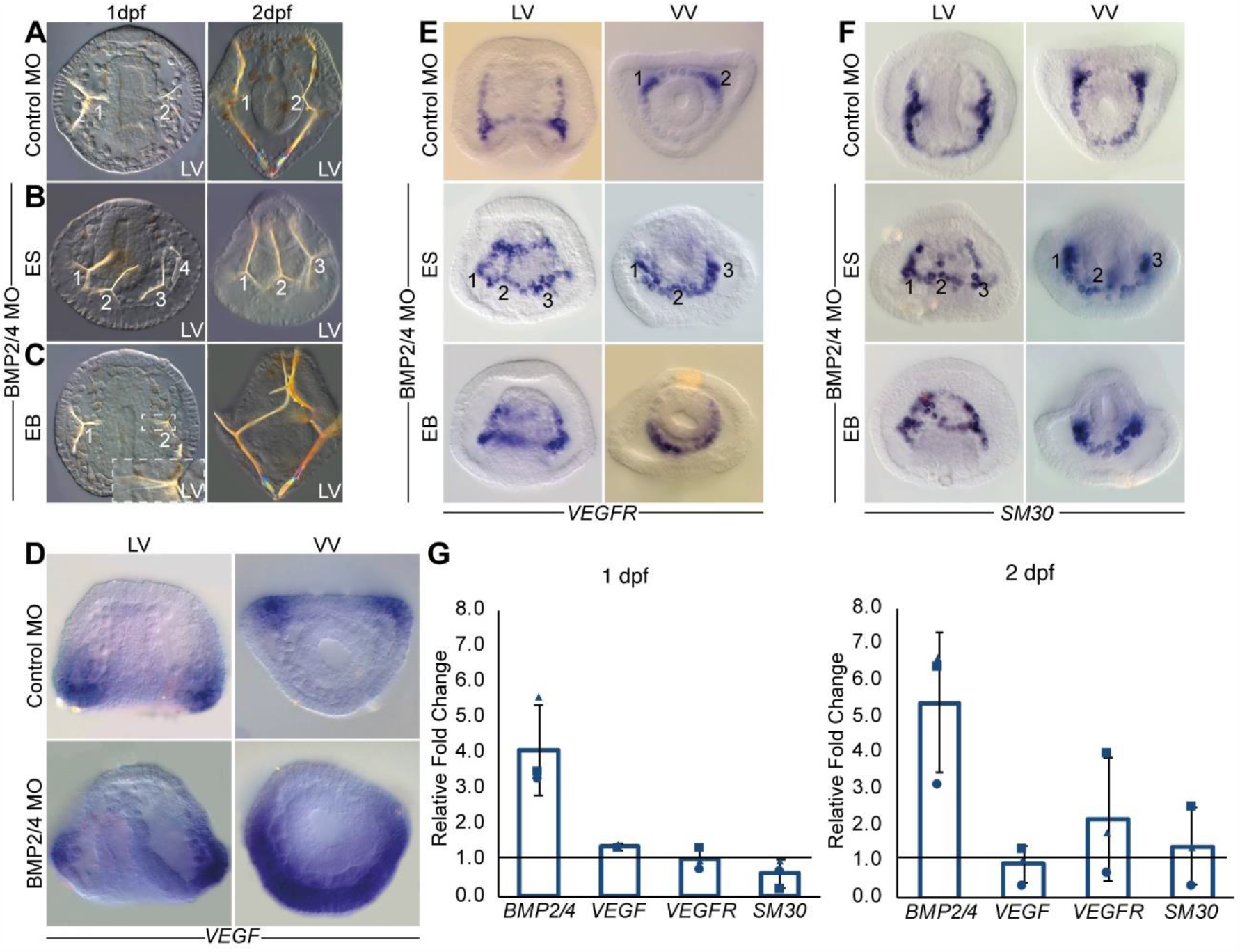
BMP2/4 controls skeletal patterning and VEGF expression. **(A)** Embryos injected with control MO show normal two spicules at 1dpf (left, 110/110 of scored embryos show this phenotype) and 2dpf (right, 56/56). **(B-C)** BMP2/4 MO injected embryos show either ectopic spicules indicated by numbers (ES, 89/169 1dpf, 120/135 2dpf) or ectopic spicule branching (EB, 39/169 at 1dpf, 15/135 at 2dpf). **(D)** VEGF expression is localized in two lateral patches in control embryo (top) and is strongly expanded in embryos injected with BMP2/4 MO at 1dpf (bottom). **(E-F)** VEGFR and SM30 expression in control embryo (top) and in BMP2/4 morphants (middle and bottom) at 1dpf. BMP2/4 MO leads to the expansion of the expression either into ectopic skeletal cell-clusters indicated by numbers (ES) or to continuous expansion (EB). LV, lateral view, VV, ventral view. Phenotypes are based on n≥3 independent biological replicates and spatial expression were observed in two independent biological replicates where n≥30 embryos were scored in each condition. **(G)** Ratio between gene expression in BMP2/4 MO compared to control MO embryos at 1dpf (left graph) and 2dpf (right graph). Bars show averages and markers indicate individual measurements of three independent biological replicates. Line marks ratio of 1 that indicates that the expression of the gene is unaffected by the perturbation. Error bars indicate standard deviation.

To further understand the regulatory links between the ectoderm and the skeletogenic GRNs, we studied the effect of BMP2/4 KD on the spatial expression of *VEGFR* and its target gene, the Spicule-Matrix protein 30 (SM30), at the gastrula stage (Duloquin et al., 2007; Morgulis et al., 2019). In control embryos, the expression of *VEGFR* is localized to the two skeletogenic cell clusters where the spicules first form (Fig. 2E) and the expression of *SM30* is noticeably enhanced in these clusters (Fig. 2F). BMP2/4 KD leads to two distinct expansion patterns of the expression of *VEGFR* and *SM30* (Fig. 2E, F). Some embryos show a continuous expansion of *SM30* and *VEGFR* expression throughout the dorsal skeletogenic cells which could drive the ectopic branching phenotype (EB in Fig. 2E, F). However, in some embryos *VEGFR* and *SM30* are expressed in three or four distinct cell clusters, which could be the cell clusters where ectopic spicules form in BMP2/4 KD (ES in Fig. 2E, F). The levels of *VEGFR* and *SM30* mRNA do not show significant change in BMP2/4 MO at one and two days post fertilization (dpf, Fig. 2G). Overall, the expression of *VEGFR* and *SM30* expands in BMP KD, which could underlie the growth of ectopic spicules and ectopic spicule branches in this condition.

The expansion of *VEGFR* and *SM30* expression in BMP morphants is probably due to the combination of direct and indirect regulation of these genes by BMP signaling. *VEGFR* and *SM30* could be directly repressed by the BMP pathway through the phosphorylation of the transcription factor SMAD1/5/8 in the dorsal skeletogenic cells. Phosphorylated SMAD1/5/8 (pSMAD1/5/8) is indeed detected in the dorsal skeletogenic cells at the gastrula stage (Lapraz et al., 2009; Lapraz et al., 2006; Luo and Su, 2012) (Fig. 1D), where it activates the expression of *tbx2/3* and *gatac* (*Duboc et al*., *2010*). The expression of *VEGFR* and *SM30* in BMP morphoants could be also enhanced indirectly, through the expansion of *VEGF* expression in these embryos (Fig. 2D). Together, our results suggest that BMP2/4 signaling controls sea urchin skeletal patterning, through the repression of *VEGF* expression in the dorsal ectoderm, and the repression of *VEGFR* and *SM30* in the dorsal skeletogenic cells.

### HIF1α does not regulate skeletal patterning and VEGF expression in the sea urchin embryo

HIF1 is one of the most potent factors in the hypoxia pathway and specifically, it activates *VEGF* expression during hypoxia induced vascularization in vertebrates (Carmeliet, 2005; Pagès and Pouysségur, 2005). Since the sea urchin HIF1α was shown to participate in early DV specification (Ben-Tabou de-Leon et al., 2013; Chang et al., 2017), we wanted to study the effect of the perturbation of this gene on sea urchin *VEGF* expression. In the sea urchin species, *Strongylocentrotus purpuratus* (*S. purpuratus*), HIF1α KD reduced the early expression of the dorsal transcription factors, Tbx2/3 and Dlx, reduced the extension of the dorsal apex and mildly reduced the elongation of the dorsal skeletal rods (Ben-Tabou de-Leon et al., 2013; Chang et al., 2017). To study the effect of HIF1α perturbation on *VEGF* expression we injected HIF1α translation MO into the eggs of the sea urchin, *P. lividus* (Fig. 3). HIF1α KD did not result with distinct skeletogenic phenotypes, in agreement with its weak effect on *S. purpuratus* skeletogenesis (Chang et al., 2017) (Fig. 3A).

**Figure 3.**
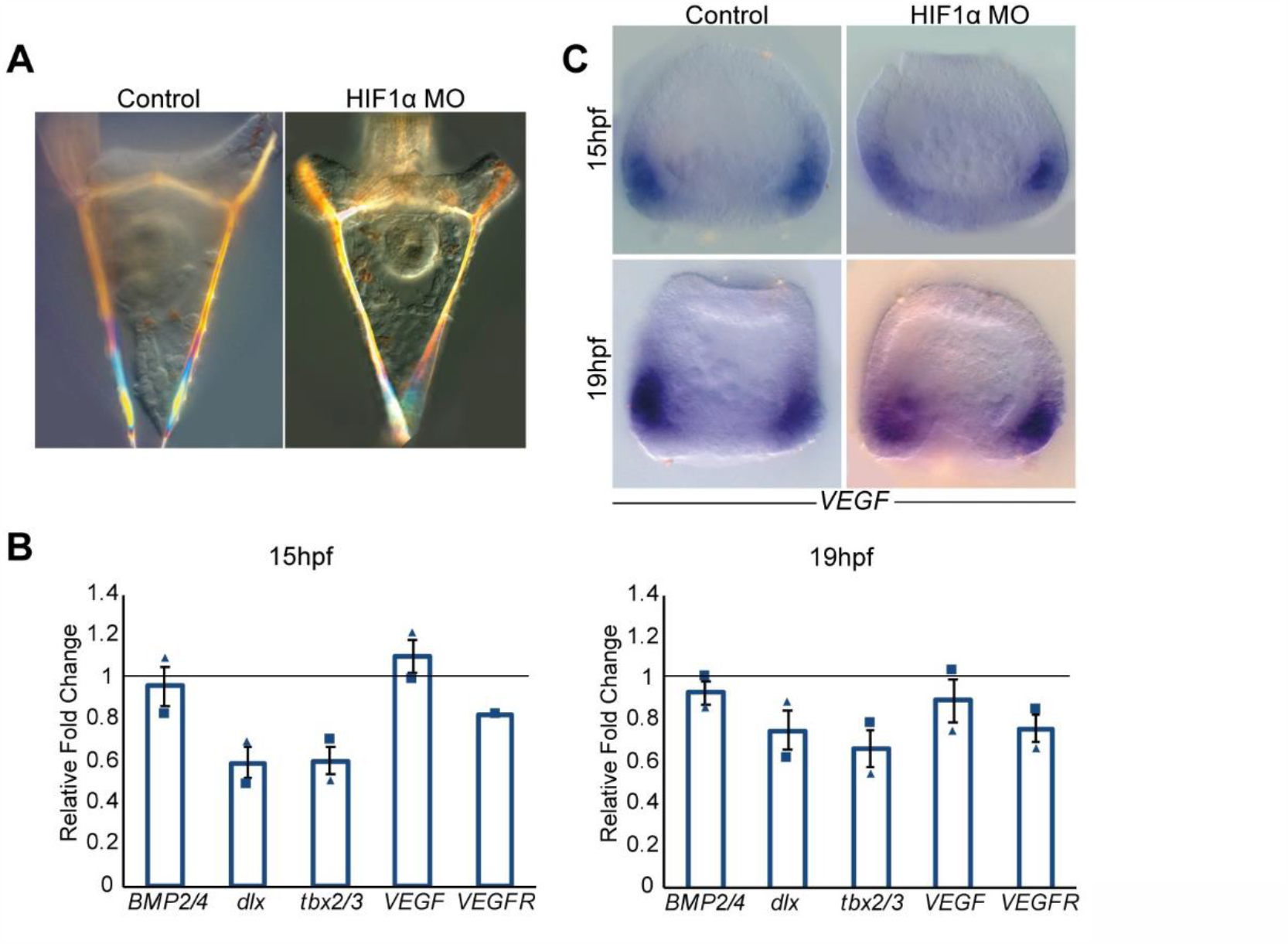
Sea urchin HIF1α does not affect skeletal patterning and VEGF expression. **(A)** Control (left) and HIF1α MO injected embryos (right) show comparable skeletal structure at 2dpf. **(B)** Ratio between gene expression in HIF1α MO compared to control MO embryos at 15hpf (left graph) and 19hpf (right graph). Bars show averages and markers indicate individual measurements of two independent biological replicates. Line marks ratio of 1 that indicates that the expression of the gene is unaffected by the perturbation. Error bars indicate standard deviation. **(C)** *VEGF* expression is similar in embryos injected with control MO (left) and HIF1α MO (right) at 15hpf (top) and 19hpf (bottom). Spatial expression were observed in two independent biological replicates where n≥30 embryos were scored in each condition.

We tested the effect of HIF1α KD on gene expression level at two developmental time points: 15 hours post-fertilization (hpf) which is equivalent to the developmental time where HIF1α activates its dorsal target genes in *S. purpuratus*, and 19hpf, when the effect of HIF1α perturbation starts to decrease in *S. purpuratus* (Ben-Tabou de-Leon et al., 2013). HIF1α KD decreases the expression level of its known target genes, *Pl*-*tbx2/3* and *Pl*-*dlx*, with a stronger reduction in the earlier time point, similarly to its effect in *S. purpuratus* (Ben-Tabou de-Leon et al., 2013), supporting the specificity of HIF1α MO (Fig. 3B). However, HIF1α KD does not affect *VEGF, VEGFR* and *BMP2/4* expression level at both times. Additionally, HIF1α KD does not affect the spatial expression of *VEGF* in the two time points (Fig. 3C). Thus, our results indicate that the role of HIF1α is restricted to dorsal ectoderm regulation, and does not interfere with skeletal patterning and VEGF regulation in the sea urchin embryo.

### Rationale of acute early and late hypoxia treatments

We sought to study the effect of transient acute hypoxia on sea urchin skeletogenesis and gene expression under hypoxic conditions that are relevant to oxygen environmental levels. The sensitivity to hypoxia changes significantly between different species and for adult sea urchin the reported sub-lethal threshold for hypoxia is 1.22 mg/L O_2_ (Sub-lethal threshold means that the animals survive this stress but their growth, reproduction and physiology are damaged (Vaquer-Sunyer and Duarte, 2008)). Water-quality surveys on sites where a massive mortality event occurred, detected levels of 0.5mg/L O_2_ and below in the seabed in depth of 10 meters and under (Altieri et al., 2017). We therefore studied the effect of growth in 0.4-0.5 mg/L O_2_, which is severe hypoxic conditions, at 18°C, that is the typical temperature for the upper water column in the Mediterranean sea (Mavropoulou A.M, 2020).

We specifically wanted to distinguish between the effect of hypoxia applied during the formation of the DV axis and hypoxia applied after the DV axis is established (Duboc et al., 2004; Lapraz et al., 2009; Nam et al., 2007; Range et al., 2007). Starting at the early blastula, the expression of Nodal, is maintained by an auto-regulation, where Nodal signaling activates the expression of the *nodal* gene (Duboc et al., 2004; Lapraz et al., 2009; Nam et al., 2007; Range et al., 2007). This could indicate that this later phase of development is less sensitive to exogenous hypoxia (Fig. 1B).

Early blastula occurs in *P. lividus* embryos under normal conditions at about 10hpf (Duboc et al., 2004; Lapraz et al., 2009), but when the embryos are grown in hypoxic conditions their development is slower and they reach this stage at 16hpf. We therefore studied the effect of growth in hypoxic conditions (0.4-0.5 mg/L O_2_) for 16 hours, from fertilization and on (early hypoxia, Figs. 4-5), and from early blastula stage and on (late hypoxia, Fig. 6). We observed significant differences in the skeletogenic phenotypes and in gene expression between these two treatments (see methods for the exact protocol).

**Figure 4.**
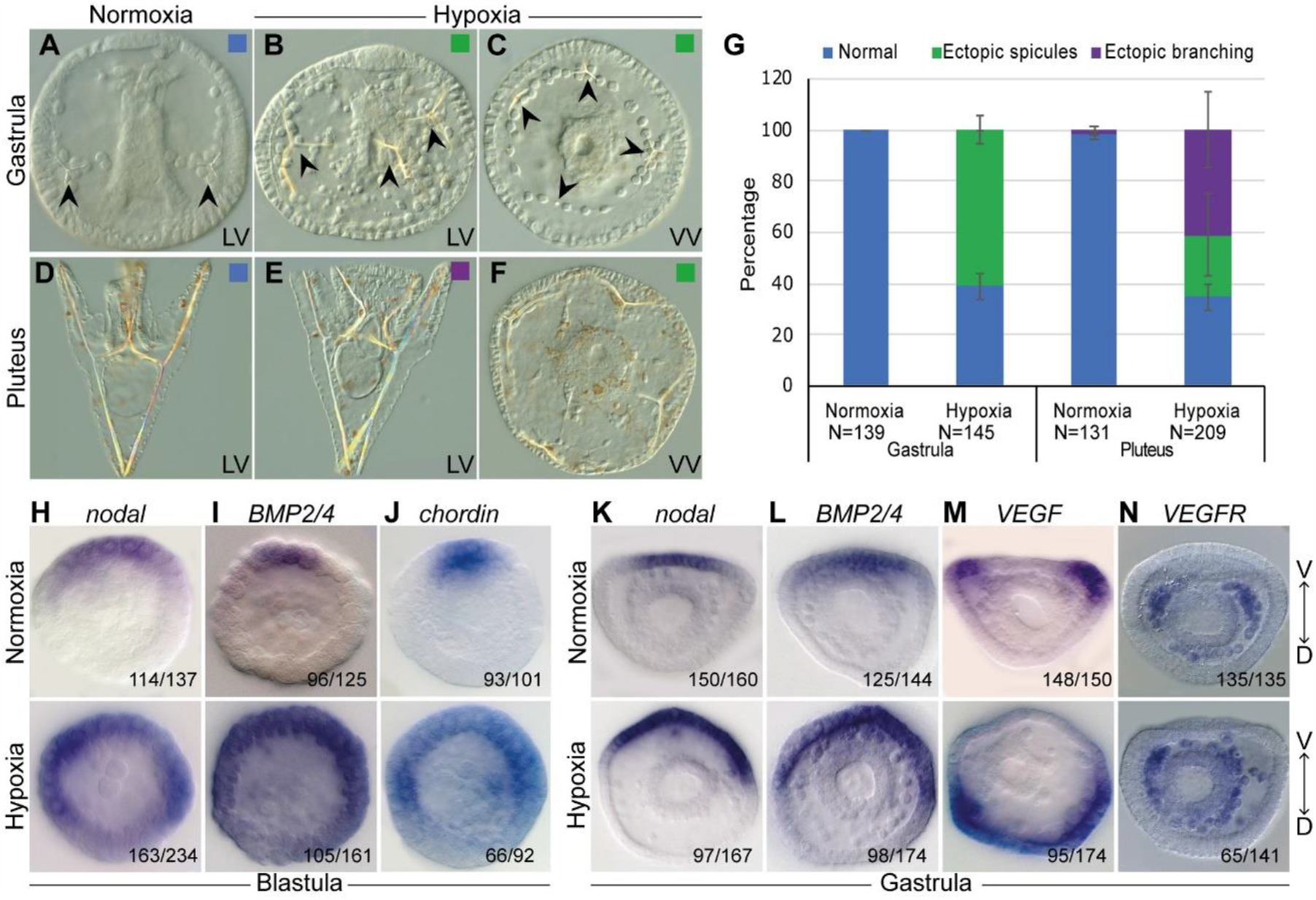
Growth in hypoxic condition leads to skeletal defects and perturbs the expression of DV and skeletal patterning genes. **(A-C)** Representative images of embryos at gastrula stage. **(A)** Embryo grown in normoxic conditions shows normal development of two spicules. **(B-C)** embryos grown in hypoxic condition show ectopic spicules, indicated by arrowheads. **(D-F)** Representative images of embryos at pluteus stage. **(D)** Embryo grown in normoxic conditions shows normal skeleton. **(E)** Embryo grown in hypoxic condition shows a normal DV axis and ectopic spicule branches. **(F)** Radialized embryo grown in hypoxic conditions that displays multiple ectopic spicules. LV, lateral view; VV, ventral view. **(G)** Quantification of skeletogenic phenotypes at gastrula stage and pluteus stage. Color code is indicated in the representative images. Error bars indicates standard deviation of three independent biological replicates. **(H-J)** Spatial expression of *nodal, BMP/4* and *chordin* genes in normoxic (top) and hypoxic embryos (bottom) at blastula stage. **(K-N)** Spatial expression of *nodal, BMP2/4, VEGF* and *VEGFR* genes in normoxic (top) and hypoxic embryos (bottom) at the gastrula stage. Embryos are presented in ventral view and the axis is labeled as V, ventral and D. Throughout H-N, the numbers at the bottom right indicate the number of embryos that show this expression pattern out of all embryos scored, based on three independent biological replicates.

**Figure 5.**
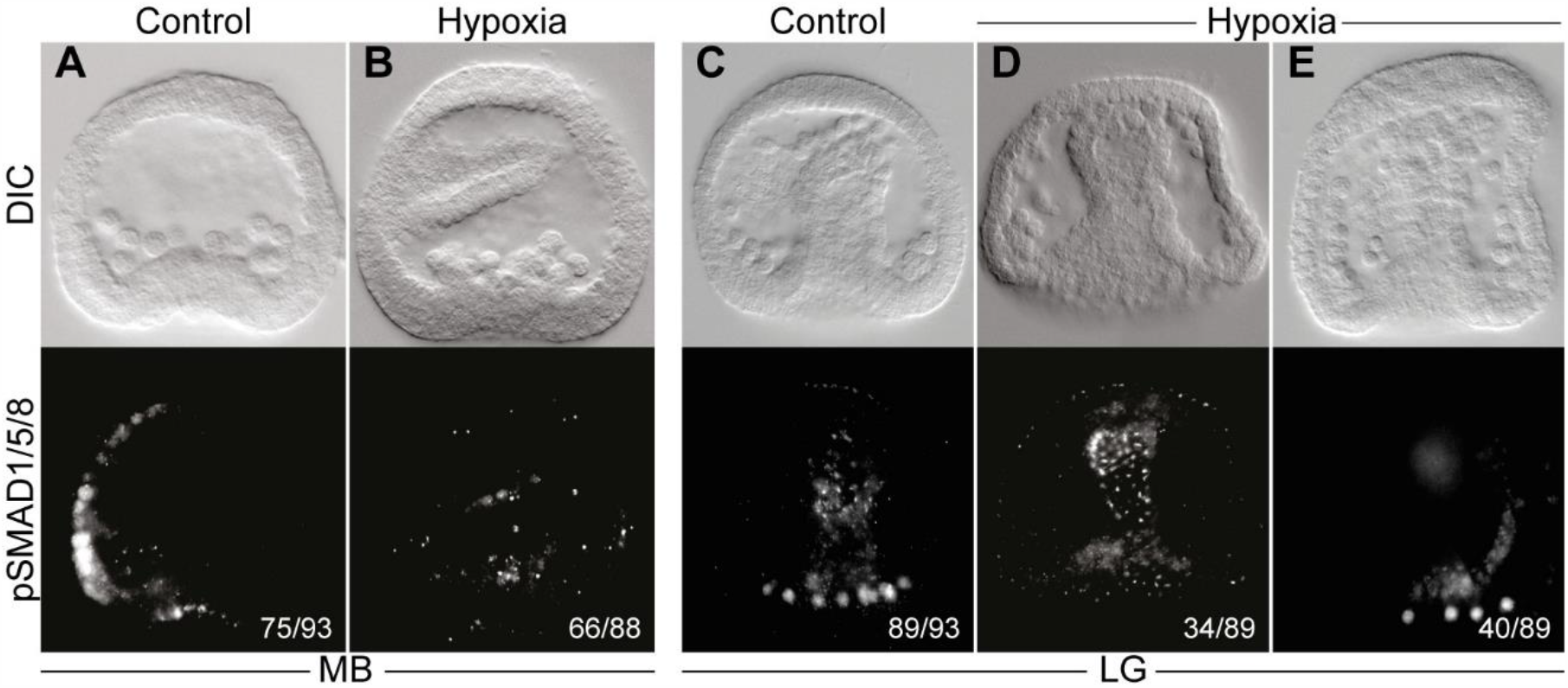
BMP activity is reduced in hypoxic conditions. **(A-B)** Nuclear pSMAD1/5/8 patterning in normoxic and hypoxic conditions at mesenchyme blastula (MB) stage. In normoxic conditions, pSMAD1/5/8 staining is detected in the dorsal ectoderm (A), while in hypoxic embryos the signal is completely abolished (B). **(C-E)** pSMAD1/5/8 staining in normoxic vs. hypoxic embryos at late gastrula (LG) stage. pSMAD1/5/8 is detected in the nuclei of the dorsal skeletogenic cells of normoxic embryos (C), while in hypoxic conditions the signal is either not detectable (D) or strongly reduced (E). DIC images of the embryos are presented in the upper row of each panel, and immunostaining of pSMAD1/5/8 of the embryos are presented in the lower row. All embryos are presented in lateral view (LV). The numbers shown on the bottom right of each figure indicate the number of embryos that show this expression pattern out of all embryos scored, based on three independent biological replicates.

**Figure 6.**
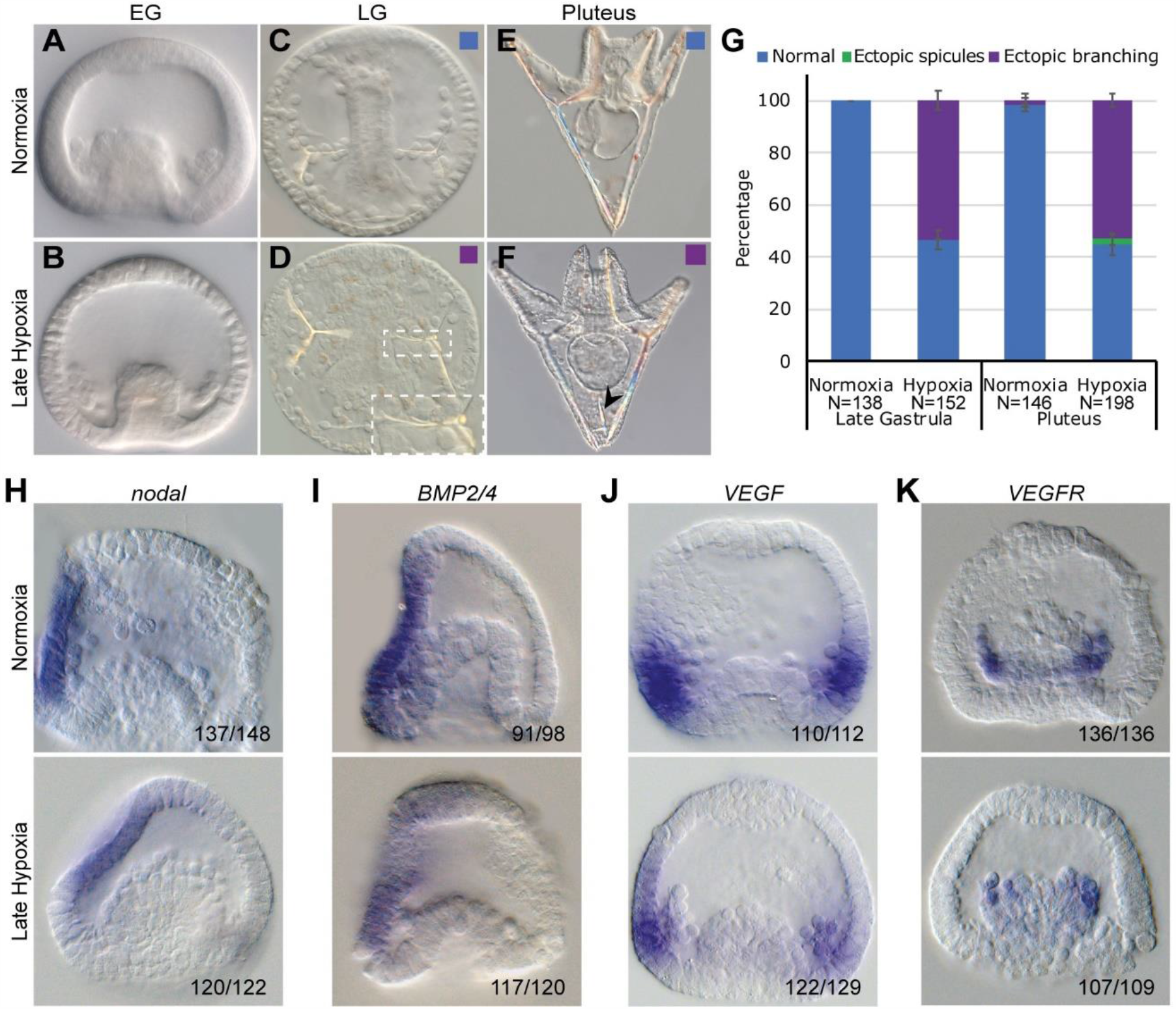
Late hypoxia affects skeletal structure but not skeletal patterning. **(A-F)** Representative images of live embryos, normoxic embryos are presented in the upper row, and equivalently staged hypoxic embryos are on the bottom. **(A-B)** Embryos at early gastrula stage show similar morphology in normoxia and hypoxia. **(C-D)** Hypoxic embryo at late gastrula stage shows two spicules with ectopic spicule branching (D) that are not observed in the normoxic embryo (C). Dashed white square is an enlarged image of the abnormal spicules. **(E-F)** Embryos at pluteus stage. Arrowhead in F, indicates an abnormal spicule growth in the hypoxic embryo. **(G)** Quantification of late hypoxia experiment over three biological replicates. Color code is indicated in the representative images. Error bars indicates standard deviation of three independent biological repeats. **(H-K)** WMISH results of *nodal, BMP2/4, VEGF* and *VEGFR* at early gastrula stage. Normoxic embryo is presented in the top and hypoxic embryo is in the bottom of each panel. On the bottom right of each figure we indicate the number of embryos that show this expression pattern out of all embryos scored, based on three independent biological replicates.

### Early hypoxia distorts skeletal patterning and expands ventral and skeletal gene expression

Embryos grown for 16hpf in hypoxic conditions applied immediately from fertilization and on (early hypoxia), are viable and develop into a normally looking blastula, but show severe DV axis disruption and skeletogenic defects from the gastrula stage and on (Fig. 4A-G). This is in agreement with previous works on *S. purpuratus* and indicates that the effect of hypoxic conditions is not species specific (Agca et al., 2009; Chang et al., 2017; Coffman et al., 2009; Coffman et al., 2004; Coffman et al., 2014). At gastrula stage, most of the embryos grown in early hypoxia show irregular skeleton with several ectopic spicules (61%, Fig. 4B, C, G). At pluteus stage, the embryos show partial recovery and display two major skeletogenic phenotypes: A strong phenotype where the skeleton is radialized, the DV axis is disrupted and multiple ectopic spicules are observed (24%, Fig. 4F, G) and a weaker phenotype where the DV axis seems normal but the skeleton shows ectopic spicule branching (41%, Fig. 4E, G). The rest of the embryos developed normally. The skeletogenic phenotypes indicate that hypoxic conditions can strongly affect skeletal patterning probably through changes in skeletogenic gene expression.

Next, we investigated the effect of hypoxia on the expression of the DV patterning genes, *nodal, BMP2/4* and *chordin*, at blastula and gastrula stages in *P. lividus*. Growth in hypoxic conditions significantly expands *nodal* spatial expression throughout the ectoderm at blastula stage, compared to the ventral localized expression of this gene in normal development (Fig. 4H), in agreement with previous studies in *S. purpuratus* (Coffman et al., 2014). The spatial expression of *BMP2/4* and *chordin* show similar expansion at this time, as expected from downstream target genes of Nodal signaling (Fig. 4I, J). At early gastrula stage, the expression of *nodal* and *BMP2/4* is expanded in embryos grown in hypoxic conditions compared to the expression of these genes in embryos grown in normoxic conditions (Fig. 4K, L). However, the expansion at gastrula stage is not throughout the ectoderm like in the blastula stage, but seems more localized to about a half of the ectoderm, in agreement with the partial phenotypic recovery at the pluteus stage (Fig. 4G).

These results suggest that hypoxia leads to the expansion of the ventral ectoderm and probably to the decrease in the dorsal ectoderm domain, which may affect the expression of key skeletogenic regulators, such as *VEGF* and *VEGFR*. Indeed, growth in hypoxic conditions shifts and expands the spatial expression of *VEGF* to one side of the ectoderm, which is most likely the dorsal ectoderm (Fig. 4M). In addition, the expression of *VEGFR* expands beyond the two lateral skeletogenic cell clusters in which it is normally localized (Fig. 4N). Furthermore, the *VEGFR* expressing cells demonstrate the perturbed migration of the skeletogenic cells in hypoxic embryos. This phenotype could be due to the expanded expression of the VEGF ligand that directs the migration of the skeletogenic cells in normal embryos. In sum, growth in hypoxic conditions perturbs the spatial organization of the skeletogenic cells and expands the ectodermal expression of *Nodal, BMP2/4, chordin* and *VEGF* and the skeletogenic expression of *VEGFR*.

### Early hypoxia reduces BMP activity which explains *VEGF* and *VEGFR* expansion

The expansion of the ventral side in hypoxic conditions suggests that BMP activity at the dorsal side might be reduced, and the reduction of the repressing BMP activity could explain *VEGF* and *VEGFR* expansion to the dorsal side. To test this hypothesis and monitor BMP activity in normal vs. hypoxic conditions, we performed immunostaining against pSMAD1/5/8. We studied pSMAD1/5/8 signal at two different developmental stages; mesenchyme blastula, when BMP activity is localized at the dorsal ectoderm (Fig. 5A), and at late gastrula, when BMP activity is localized at the dorsal skeletogenic cells (Fig. 5C). Hypoxic conditions completely abolish pSMAD1/5/8 signal from the nuclei of the dorsal ectodermal cells at mesenchyme blastula stage (Fig. 5B). At late gastrula stage, hypoxic conditions eliminate the pSMAD1/5/8 signal from the dorsal skeletogenic cells (Fig. 5D), or strongly reduce it (Fig. 5E). These results indicate, that despite *BMP2/4* expansion in hypoxic embryos, its activity is reduced during hypoxia. The reduced activity can be explained by the expansion of BMP antagonist, Chordin, during hypoxic conditions (Fig. 5C). Together, these results show that BMP activity in the dorsal ectoderm and in the dorsal skeletogenic cells is reduced in hypoxic conditions. Apparently, the reduction of BMP activity removes the repression of *VEGF* and *VEGFR* at the dorsal embryonic domains, leads to their expansion to this domain and to the disruption of skeletal patterning.

### Late hypoxia mildly affects skeletogenesis and doesn’t affect DV and skeletal regulatory genes

Our studies show that early hypoxia strongly affects the spatial activity of the main regulators of DV axis formation, Nodal and BMP2/4, and the perturbation of these factors affects skeletal patterning and *VEGF, VEGFR* and SM30 expression. Next, we wanted to test whether hypoxia affects skeletogenesis after the DV axis is formed and to investigate the effect of late hypoxic conditions on regulatory gene expression. Thus, we studied the skeletogenic phenotypes of hypoxia applied between 10hpf and 26hpf, which is after the DV axis is established, as explained above. Embryos grown in late hypoxia showed a delayed development and at 26hpf were equivalent to early gastrula stage in normoxic embryos (Fig. 6A, B). At late gastrula and pluteus stages, almost all the embryos grown in late hypoxia show normal skeletal patterning with the two spicules correctly positioned at the two lateral sides (Fig. 6C-G). More than half of the embryos grown in late hypoxia developed ectopic skeletal branching in these two stages, and at pluteus stage, about 2% of the embryos show radialized skeleton with ectopic spicules. Overall, late hypoxia induces skeletal defects, such as ectopic branching, but it hardly affects skeletal patterning.

We next studied the effect of late hypoxic conditions on the expression of the key regulatory genes investigated above. Late hypoxia treatment does not affect the spatial expression of *nodal* (Fig. 6H), in agreement with the normal formation of the DV axis and normal skeletal patterning in this condition. Furthermore, late hypoxia does not affect the spatial expression pattern of *BMP2/4, VEGF* and *VEGFR* genes, so these genes are probably not the mediators of the observed mild skeletal defects (Fig. 6I-K). Thus, after the DV axis forms, the expression of the upstream patterning and skeletogenesis regulators, *nodal, BMP2/4, VEGF* and *VEGFR* is not affected by hypoxic conditions and the skeletal patterning is overall normal.

## Discussion

GRNs are the genomically encoded programs that control embryonic development, but the environmental conditions in which these GRNs operate can significantly affect their outcome (Beldade et al., 2011; Smith et al., 2018). Particularly, the use of hypoxia and redox gradients to control developmental processes in various phyla, might make the embryos more sensitive to low oxygen levels that are becoming more common in the ocean (Compernolle et al., 2003; Cordeiro and Tanaka, 2020; Dunwoodie, 2009; Semenza, 2012). The structure of the developmental GRN defines its function during environmental hypoxia and underlies the response and resilience to hypoxia during embryogenesis. Here we studied the regulatory linkages and response to transient acute hypoxia of the GRNs that control DV patterning and skeletogenesis in the sea urchin embryo (Fig. 7A, B). We discovered that hypoxia applied during the time where a redox gradient guides the DV axis formation results with a major disruption of the spatial expression of key regulatory genes which explains the radial skeleton formation in these embryos. However, once the DV axis is established, these regulatory genes are no longer affected by hypoxic conditions and skeletal patterning is largely normal. While this suggests that embryos could overcome transient hypoxia if it occurs after their DV axis is formed, hypoxic conditions in natural habitats can last for days and weeks, which is much more than the embryos can tolerate (Altieri et al., 2017). Below we discuss our main findings and their possible implications.

**Figure 7.**
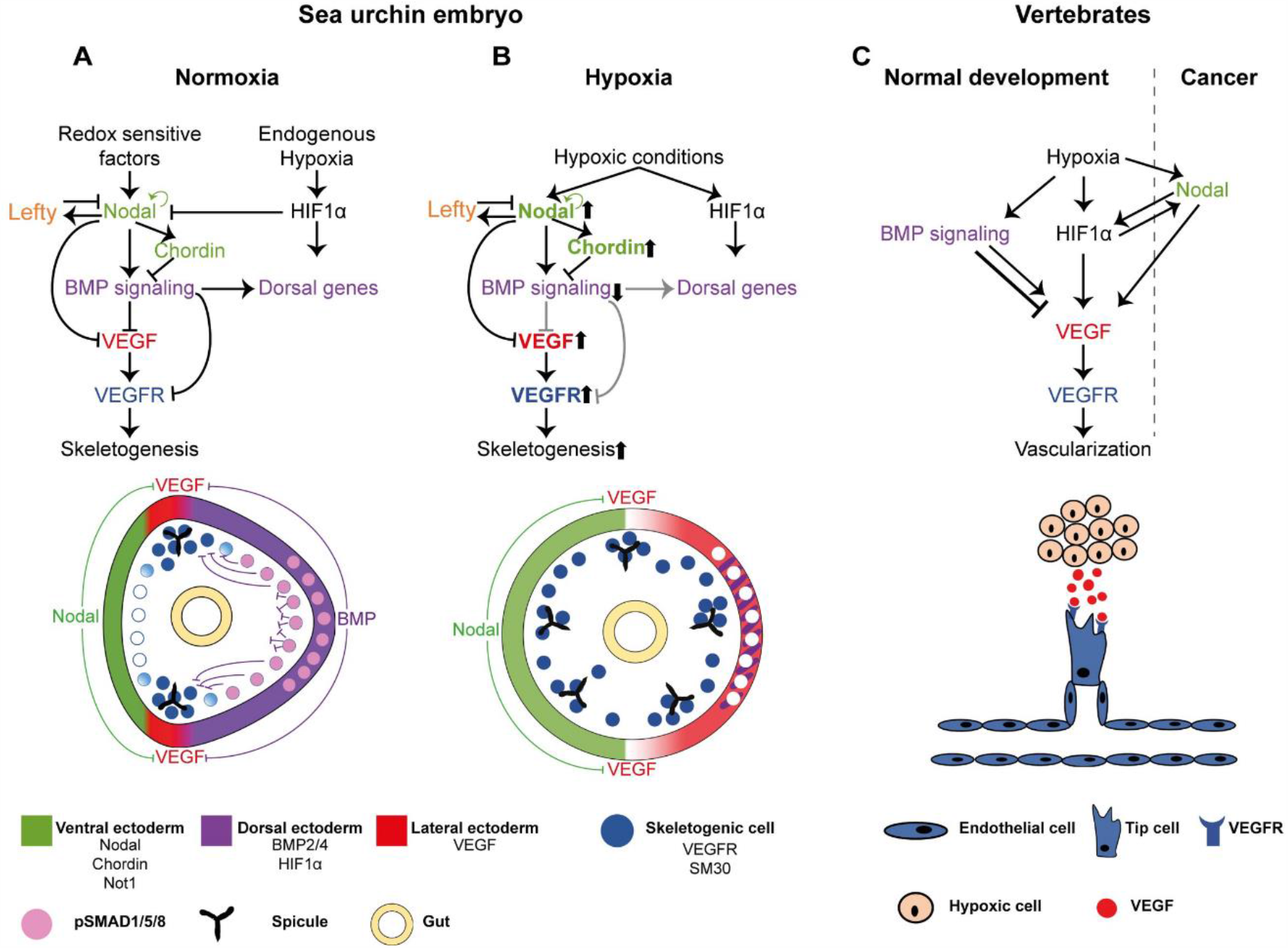
The interactions between the DV and skeletogenic GRNs, the response to early hypoxia and the similarities to the regulation of vertebrate’s vascularization. **(A-B)** Diagrams showing our proposed model for skeletal patterning in normal conditions **(A)** and Hypoxic conditions **(B)**. Color codes are indicated in the bottom part of the figure. **(A)** The regulatory interactions between Nodal, BMP, HIF1α and VEGF signaling during normal development. BMP represses *VEGF, VEGFR* and *SM30* expression in the dorsal side and HIF1α does not regulate VEGF expression in the sea urchin embryo **(B)** The modification of the regulatory states in hypoxic conditions applied at early development, revealed in this work. Early hypoxia expands *nodal* expression and reduces BMP activity and the dorsal ectoderm. The reduction of BMP activity leads to an expansion of *VEGF*, VEGFR and *SM30* expression in the dorsal side and growth of ectopic skeletal centers. Ascending arrows near a gene name indicate enhanced activity, while descending arrows indicate reduced activity. Gray regulatory links indicate inactive connections under hypoxic conditions. **(C)** Diagram showing the relevant regulatory interactions during vertebrates’ vascularization in normal development and in cancer, see text for explanations.

Our findings illuminate the regulatory interactions between the DV and skeletogenic GRNs that underlie skeletal patterning in the sea urchin embryo. Previous studies had shown that *VEGF* expression is restricted from the ventral ectoderm by Nodal’s target, Not1 (Li et al., 2012), but the mechanism that excludes *VEGF* expression from the dorsal ectoderm was not known. Here we show that BMP activity excludes both *VEGF* from the dorsal ectoderm and *VEGFR* and *SM30* from the dorsal skeletogenic cells, and this exclusion is necessary for spicule initiation to occur only in the ventro-lateral skeletogenic clusters (Fig. 2). We also show that HIF1α, a key activator of VEGF in vertebrates’ vascularization (Dunwoodie, 2009; Pagès and Pouysségur, 2005) does not regulate VEGF signaling during early sea urchin development (Fig. 3, 7A). Apparently, the regulatory function of this factor in normal sea urchin development is limited to shaping *nodal* expression domain in the early blastula (Chang et al., 2017), and to activating early dorsal gene expression (Ben-Tabou de-Leon et al., 2013). Thus, BMP signaling restricts VEGF activity to the ventro-lateral skeletogenic clusters and this restriction is required for the exclusion of spicule formation from the dorsal skeletogenic cells in normal sea urchin embryos (Fig. 7A).

Early hypoxia in sea urchin embryos strongly distorts the spatial expression of DV and skeletogenic patterning genes, which leads to the formation of ectopic spicules and embryo radialization (Fig. 4, 7B). Previous studies have shown that hypoxic embryos are ventralized (Agca et al., 2009) and that *nodal* expression expands in hypoxic conditions (Coffman et al., 2014). Here we revealed the cascade of regulatory interactions that underlie embryo ventralization and the formation of ectopic spicules. Early hypoxia leads to the expansion of *nodal* to the dorsal side, which leads to the expansion of its targets, *BMP2/4* and *chordin* in this condition (Fig. 4H-L, Fig. 7B). The activity of BMP signaling is significantly reduced in both the dorsal ectoderm and dorsal skeletogenic cells as evident from pSMAD1/5/8 staining (Fig. 5). This reduction is probably due to the expansion of the expression of BMP antagonist, *chordin*, into the dorsal side, which blocks BMP activity in early hypoxia embryos (Fig. 4J, 7B). These changes in the spatial activity of Nodal and BMP signaling drive the shift and expansion of *VEGF* and *VEGFR* expression to the dorsal ectoderm and dorsal skeletogenic cells, respectively (Fig. 4M, N 7B). The expansion of VEGF activity to the dorsal skeletogenic cells explains the formation of ectopic spicules in the dorsal side in early hypoxic condition (Fig. 7B). Hence, early hypoxia expands *nodal* expression which reduces BMP activity and specifically, removes the dorsal repression of VEGF signaling, which leads to the formation of ectopic spicules in the dorsal side.

In striking difference to the strong effect of early hypoxia on the expression of DV and skeletogenic regulatory genes, hypoxia applied after the DV axis is establish does not affect the expression of these genes and results with overall normal skeletogenic patterning (Fig. 6). This resilience of the DV GRN to late hypoxia can be explained by the structure of the GRN, that includes positive and negative feedback loops that restrain Nodal activity (Fig. 7A,B, (Duboc et al., 2008; Nam et al., 2007; Range et al., 2007)). At the early blastula stage, *nodal* expression is maintained by the Nodal pathway through the transcription factor SMAD2/3 (Nam et al., 2007; Range et al., 2007), and *nodal* spatial expression is restricted by its antagonist, Lefty, that is also activated by the Nodal pathway (Fig. 7A, (Duboc et al., 2008)). Once the spatial domain of Nodal activity is established and stabilized by the Nodal-Lefty feedback loops, it is not disrupted by hypoxic conditions, indicating that the redox state is no longer a factor in *nodal* regulation at this stage (Fig. 6H). Nodal spatial activity defines the domain of BMP activity through the Nodal-BMP2/4-Chordin incoherent feedforward loop, which restricts VEGF activity that leads to normal skeletogenic patterning, in late hypoxia embryos (Figs. 6, 7A). This structure of the DV GRN could also underlie the relative restriction of *nodal* expression at the gastrula stage compared to its broad expression at the blastula stage (Fig. 4H, K) and the partial recovery of skeletal patterning in the pluteus stage in early hypoxia (Fig. 4G). Overall, the structure of the DV GRN enables it to partially recover the effect of early hypoxia at later developmental stages and makes it resilient to hypoxia applied after the DV axis had formed.

While the effect of the hypoxic conditions on the regulatory cascade downstream of Nodal signaling is quite clear from our findings, the cause of nodal expansion in early hypoxia requires further investigation. Early hypoxia was shown to expand the expression of the HIF1α protein that is normally localized at the dorsal side (Chang et al., 2017). HIF1α transiently represses early *nodal* expression (Chang et al., 2017), yet, HIF1α expansion does not restrict *nodal* expression that expands dorsally in early hypoxia (Fig. 4). Hypoxia was shown to increase ROS levels in cancer cells, smooth muscle cells and endothelial cells, apparently due to its effect on the mitochondria electron transport chain (Chi et al., 2010; Desireddi et al., 2010; Fuhrmann and Brune, 2017; Medini et al., 2020; Tafani et al., 2016). If hypoxia affects the mitochondria in the sea urchin embryo and further increases the ROS levels at the already oxidizing side (Fig. 1), this could underlie the expansion of *nodal* expression in early hypoxia. Thus, *nodal* expansion could be the result of the change in the redox state in hypoxic conditions and the effect of this change on the redox sensitive transcription factors that control *nodal* expression (Agca et al., 2009; Coffman et al., 2004; Coffman et al., 2014; Range et al., 2007). Future research will hopefully illuminate this intriguing regulatory mechanism.

Our findings illuminate some similarities between the GRNs that pattern the DV axis and skeletogenesis in the sea urchin embryo and the upstream regulation of vertebrate’s vascularization (Lee et al., 2009; Ushio-Fukai and Nakamura, 2008). Hypoxia and redox gradients that regulate DV axis formation and skeletal patterning in the sea urchin embryo, were shown to induce angiogenesis in vertebrates during normal development and in cancer (Chi et al., 2010; Potente et al., 2011). The regulatory interactions between BMP and VEGF that are essential for sea urchin skeletal patterning, also control vertebrates’ vascularization, however, they are rather complex: BMP activates VEGF and induces vascularization in some tissues, while it represses VEGF in other tissues (Bai et al., 2013; Dyer et al., 2014; Garcia de Vinuesa et al., 2016; He and Chen, 2005; Wiley et al., 2011). The Nodal pathway does not participate in hypoxia induced vascularization during normal development in vertebrates, however, in various cancer cells, hypoxia drives Nodal expression, which then promotes *VEGF* expression and angiogenesis (Fig. 7C, (Hueng et al., 2011; Quail et al., 2011; Quail et al., 2012)). The transcription factor HIF1α is a key activator of VEGF expression and angiogenesis in vertebrates, but the sea urchin HIF1α does not regulate VEGF signaling during normal development. Sea urchin HIF1α activity is limited to the transient inhibition of *nodal* and the early activation of dorsal genes (Fig. 3, 7) (Ben-Tabou de-Leon et al., 2013; Chang et al., 2017). Overall, regulatory interactions between Nodal, BMP, HIF1 and VEGF pathways and their modulation by hypoxic conditions are observed both during DV and skeletal patterning in the sea urchin embryo and in vertebrates’ vascularization, but there are some apparent differences in the linkages. The participation of these common pathways together with the similarity between the skeletogenic and the vascularization GRNs (Morgulis et al., 2019; Oliveri et al., 2008) might indicate that these upstream patterning programs diverged from a common ancestral GRN; yet we cannot exclude convergent evolution at this stage.

Our findings have implications on the effect of ocean deoxygenation on embryos that use hypoxia and redox signaling in their development, yet, the major differences between lab experiments and field conditions should be considered. Our analyses and previous studies suggest that the use of hypoxia and redox gradients makes the sea urchin GRNs highly sensitive to acute hypoxia applied in its early developmental stages, but the GRNs are less sensitive to hypoxia applied after the establishment of the DV axis. Yet, hypoxia events in the ocean and in the coastal zones can last for weeks and their lethal effect is observed for months after (Altieri et al., 2017; Hughes et al., 2020). So even if the sea urchin embryos can survive 16 hours of hypoxia, they will probably die in longer periods of low oxygen. Furthermore, in other organisms ROS and hypoxia signaling regulate multiple developmental processes and in some cases, these processes last throughout embryogenesis, which could make the embryos of these organisms even more sensitive to hypoxia than sea urchin embryos (Breus and Dickmeis, 2020; Coffman and Su, 2019; Cordeiro and Tanaka, 2020). Within these alarming notions, lab experiments can show distinct and even opposing trends then experiments that are done in the field due to the increased and unexpected complexity of natural sites (Foo et al., 2020). Therefore, further hypoxia studies guided by environmental changes should be done in the field, to elucidate the sensitivity and resilience of the molecular response to hypoxia in marine embryos in their natural habitat.

## Materials and Methods

### Animals and embryo cultures

Adult *P. lividus* sea urchins were purchased from the Institute of Oceanographic and Limnological Research (IOLR) in Eilat, Israel. Eggs and sperm were obtained by injection 0.5M KCl solution to adult sea urchins. Embryos were cultured in artificial seawater (ASW) at 18°C.

### Microinjection, RNA extraction and Reverse-transcription

The design and preparation of novel morpholino (MO) was done in genetools (http://www.gene-tools.com). Translation of *HIF1α* was blocked by the microinjection of 400-700µM *HIF1α*-MO into sea urchin eggs. *HIF1α*-MO sequence: 5’-GGTCGCCATAATCAGTCTCTGTTTC-3’. Translation of *BMP2/4* was blocked by the microinjection of 400-600µM. *BMP2/4*-MO sequence: 5’-GACCCAGTTTGAGGTGGTAACCAT-3’, this MO has been characterized in previous studies (Duboc et al., 2004). The control MO is Random commercial MO which does not have any effect on embryo development, along with 1µg/ml rhodamine dextran (D3329 Molecular probes, OR, USA) and 0.12M KCl. Total RNA was extracted from injected sea urchin embryos (≥120 injected embryos) using RNeasy Micro Kit (50) from QIAGEN (#74004) according to the kit protocol using DNase treatment from RNease-Free DNase Set-Qiagen (50) (#79254). Elution was done in 16.5µl nuclease-free ultra-pure water. Extracted RNAs were then reverse transcribed into cDNA by using SuperScript™ II Reverse Transcriptase (Thermo Fisher scientific 18064022) (10 min 25°C, 2hr in 25°C, 85°C for 5 min).

### Quantitative-PCR (qPCR) analysis

qPCR was performed using the CFX384 Touch™ Real-Time PCR Detection System #1855485. Reactions were carried out in 10μl volume including: 5µl SYBR BioRad IQ SYBR Green Supermix (#1725125), 2.5μl of 1.2μM forward and reverse gene specific primers and 2.5μl of cDNA (qPCR primers used in this study are listed in Table S1). Each cDNA sample was run in triplicate, for every candidate gene, ubiquitin was used as internal control. The reactions thermal profile was: 95°C for 3 minutes followed by 40 amplification cycles of 95°C for 10 seconds and 55°C for 30 sec. Dissociation analysis was performed at the end of each reaction to confirm the amplification specificity. Primer sets for all tested genes were designed using Primer3Plus (http://www.bioinformatics.nl/cgi-bin/primer3plus/primer3plus.cgi/). Results are presented as the means and standard error of at least two biological replicates. The comparison to an internal standard (ubiquitin) was done in order to determine the expression level of the gene, and the change in the expression levels were measured in comparison to the expression level of the gene in control MO.

### Hypoxia treatment

ASW were treated with 99.5% Nitrogen (N_2_) and 0.5% Oxygen (O_2_) to decrease the oxygen solubility in ASW till the dissolved O_2_ level was 0.4-0.5 mg/L, creating hypoxic ASW. Embryos were transferred into petri-dish that contains the hypoxic ASW, then the dishes were incubated in a hypoxia chamber at 18°C. The hypoxia chamber is a sealed box that receives a constant flow of 99.5% N_2_ and 0.5% O_2_. To distinguish between the direct effect that hypoxic conditions might have on skeletogenesis and its effect on DV patterning, we studied the skeletogenic phenotypes of hypoxia applied immediately after fertilization (early hypoxic condition) and the effect of hypoxia applied after the DV axis was established (late hypoxic condition). In early hypoxia treatment, the eggs were fertilized, their fertilization envelope was immediately removed and the zygotes were incubated in the hypoxia chamber for 16 hours. In late hypoxia treatment, the eggs were fertilized and the embryos were cultured under normoxic conditions for 10 hours until the blastula stage. Then, the embryos were transferred into the hypoxia chamber and incubated in hypoxic conditions for 16 hours. After 16 hours in hypoxic conditions the embryos were removed from the hypoxia chamber and cultured in normoxic conditions until the pluteus stage.

### Probe design and WMISH procedure

WMISH probe preparation and WMISH procedure were performed as described in (Morgulis et al., 2019). Primer list is provided in table S2.

### Removal of fertilization envelope

To perform WMISH on sea urchin embryos at early blastula stage, the fertilization envelope were removed; Fertilized eggs were incubated in presence of Paraminobenzoic acid (PABA, A6928, Sigma) and Amino triazole (ATA, A8056 Sigma) (2mM each at final concentration) to soften the fertilization envelope (FE). After microscopy visualization of FE, FE were removed by flow the zygotes through a 75µm mesh four times. Next, the embryos were washed three times with ASW and grown till the indicated collection time points.

### Immunostaining

Immunostaining of pSMAD1/5/8 antibody was done similarly to (Lapraz et al., 2009) with minor modifications. Embryos were fixed in 4% paraformaldehyde, 33mM Maleic acid buffer pH7, 166mM NaCl, for 10 minutes at room temperature, then embryos exposed to Methanol for 1 minute. Embryos were washed four times with PBST, then incubation for 1 hour in blocking solution (PBST and 4% sheep serum), followed by incubation with primary Antibody against pSMAD1/5/8 (the antibody was purchased from Cell Signaling Technology; no. 9511) at 1:200 dilution in blocking solution, overnight at 4°C. Embryos were then washed four time in PBST, then the secondary Antibody was added to the embryos (Peroxidase-conjugated AffiniPure Goat Anti-Rabbit IgG; no. 111-035-003) diluted 1:200 in blocking solution and incubated for 1 hour in room temperature, followed by four washes with PBST. Store solution (PBST in 50% glycerol) at 4°C.

### Imaging

All images presented in this study were generated on a Zeiss Axio Imager M2.

## Acknowledgments

We thank Imad Shams for generously providing the hypoxic chamber and gas controller; Eli shemesh for providing the oxygen sensor; Shlomo Ben-Tabou de-Leon for technical assistance with the hypoxic system build-up; David Ben-Ezra for his help with sea urchin handling; Miri Morgulis for helpful comments and suggestions. This work was supported by the Israel Science Foundation Grants 41/14 and 211/20 (to Smadar Ben-Tabou de-Leon).

