## Supplemental tables for "The tolerance to hypoxia is defined by a time-sensitive response of the gene regulatory network in sea urchin embryos"

**Supplementary Materials**

**Table S1:** List of qPCR primers used in this study

| Gene Name | Forward primer | Reverse primer |
|---|---|---|
| *BMP2/4* | GTACCGGTCGCATACACAAG | TGTGTCTGTGCTGCTCTGTA |
| *dlx* | GGGCATCCTCCAATTTATGA | GCAAGGTATTGGGTCTGGTG |
| *tbx2/3* | TATTACCACCTCGCCTCAGC | CCTGGATGTTCGCCGAATTT |
| *VEGF* | GCTCATGGTTCTCTCGAAGG | CCCGCTGAGATAACATTGGT |
| *VEGFR* | CACTGGGAGATATCGGTGCT | ACGGTTGCCCACGATAAATA |
| *SM30* | AGGTGGTTTCCCTGGACAAG | TTGGGCATGTCTCCTGTCTT |

**Table S2:** List of WMISH PCR primers used in this study

| Gene Name | Forward primer | Reverse primer |
|---|---|---|
| *BMP2/4* | GTGGCGAAAGAGGAGCGAC | GATGGTTCTGATCAAGAAGTC |
| *nodal* | CATCCATCGGAGCAACTCTTC | CAAAGTTCAAAATCGAATCGGC |
| *chordin* | GACCATGTACCGTGTCGTGATTTATAC | CCCTGAGCCAAACCATCACGG |
| *VEGFR* | GTAACTCATAATCCATGCTAC | GCTACTATTCAATGTCATTTC |
| *SM30* | CCTCCCCCTTTCGTGTTATAAATG | GCAAAACAACTTCTTCGTCGGTC |
